# Image Segmentation based on Relative Motion and Relative Disparity Cues in Topographically Organized Areas of Human Visual Cortex

**DOI:** 10.1101/208033

**Authors:** Peter J. Kohler, Benoit R. Cottereau, Anthony M. Norcia

**Affiliations:** Department of Psychology, Stanford University Jordan Hall, Building 420, 450 Serra Mall, Stanford, CA 94305; Université de Toulouse, Centre de Recherche Cerveau et Cognition, Toulouse, France; Centre National de la Recherche Scientifique, Toulouse Cedex, France

## Abstract

The borders between objects and their backgrounds create discontinuities in image feature maps that can be used to recover object shape. Here we used functional magnetic resonance imaging (fMRI) to study the sensitivity of visual cortex to two of the most important image segmentation cues: relative motion and relative disparity. Relative motion and disparity cues were isolated using random-dot kinematograms and stereograms, respectively. For motion-defined boundaries, we found a strong retinotopically organized representation of a 2-degree radius motion-defined disk, starting in V1 and extending though V2 and V3. In the surrounding region, we observed phase-inverted activations indicative of suppression, extending out to at least 6 degrees of retinal eccentricity. For relative disparity, figure responses were only robust in V3, while suppression was observed in all early visual areas. When attention was captured at fixation, figure responses persisted while suppression did not, suggesting that suppression is generated by attentional feedback from higher-order visual areas. Outside of the early visual areas, several areas were sensitive to both types of cues, most notably hV4, LO1 and V3B, making them additional candidate areas for motion- and disparity-cue combination. The overall pattern of extra-striate activations is consistent with recent three-stream models of cortical organization.

## Introduction

The boundaries between objects and their background give rise to discontinuities in multiple feature maps. Relative motion and relative disparity are two strongly related parallax cues, that each contribute independently to image segmentation and the perception of shape, and can be combined to disambiguate 3D object and scene structure^1–3^. Sensitivity to image discontinuities created by relative motion has been observed in both early and higher-level visual areas. Single unit responses to relative motion information are robust in primate V1 and V2^4–9^, as well as in MT^10–13^ and IT^14^, although selectivity for the orientation of a motion-defined boundary may not arise until V2^9,15^. In humans, functional magnetic resonance imaging (fMRI) studies^16,17^ identified an area in dorsomedial occipital cortex, originally termed the kinetic occipital area (KO), that was activated by relative motion in texture-defined bars. However, other human fMRI studies at that time found that this stimulus also produces activations in V1, V2, V3 and in the hMT+ complex that includes the homologue of macaque MT^18–20^. Later work made more extensive measurements in topographically organized visual areas and used fMRI adaptation to identify selectivity for the orientation of motion boundaries in areas V3A, V3B, LO1, LO2 and V7^21^, which partially overlapped with functionally-defined area KO.

Studies in macaque suggest that image discontinuities generated by relative disparity are not encoded before V2^22–24^. Sensitivity to disparity discontinuities has also been found in V3^25^, V4^26,27^, IT^28^ and, depending on the precise definition of relative disparity, in MT^29^. In humans, the functional localization of relative disparity processing is less well established. Several studies have compared fMRI BOLD responses to displays with multiple disparity planes to responses produced by a single depth-plane, and found dominant activations in V3A^30–32^. Because these studies contrasted non-zero disparity and zero disparity displays, the effects could be driven by tuning for absolute or relative disparity, or for both. This confound was addressed by Neri and colleagues^33^, who used fMRI adaptation to measure responses to both absolute and relative disparity separately. They found that areas V4 and V8 (and to a lesser degree early visual areas V1, V2 and V3) showed adaptation effects for both relative and absolute disparity, but that adaptation was only present for absolute disparity in V3A, MT and V7. A more recent study of cue combination reported reliable classification of depth defined by motion and disparity in several visual areas, but used an experimental design that allowed classification to be driven by tuning for absolute or relative signals, or for both^1^.

Here we seek to integrate this past work by measuring fMRI BOLD responses to simple figure-ground displays based on relative motion and relative disparity in 18 topographically organized visual areas. We use disk-annulus stimulus configurations that are well-matched between motion and disparity, and an experimental design that controls for the contribution of absolute signals to the measured responses. The use of a figure-ground stimulus configuration is also expected to drive global shape processes and thus activate object-sensitive cortical areas^34–36^. Finally, we conduct a control experiment where attention is captured at fixation, to investigate the influence of top-down attention on responses.

## Results

### Eccentricity analysis in early visual cortex

We begin by describing results from early visual areas V1, V2 and V3, each of which have strong retinotopic representations of the visual field. To visualize and quantify the existence and retinotopic specificity of the representations of the motion and disparity-defined figures on the cortical surface we used a retinotopy template^37^ to define eccentricity-based sub-ROIs and then used a fMRI localizer task to verify the accuracy of the template. The goal of the sub-ROI analysis was to track the responses to central (figure), boundary and outer (ground) regions of the stimulus across the surfaces of the early visual areas. The localizer task consisted of the alternation across 12 sec block of high contrast dynamic textures in the disk with high contrast dynamic textures in the background (see Fig. 1A). We computed the signed amplitude values over the period of block alternation, by applying a Fourier transform to the average time course (see Methods). The results of the contrast-based localizer condition are plotted in Figure 2A. Positive amplitudes indicate relatively stronger responses to the A block, during which the disc was presented, compared to the B block, during which the annulus was presented. Negative amplitudes indicate the opposite. For convenience, we will refer to the former as “disk responses” and the latter as “annulus responses”. The eccentricity axis was derived independently, based on the template.

**Figure 1.**
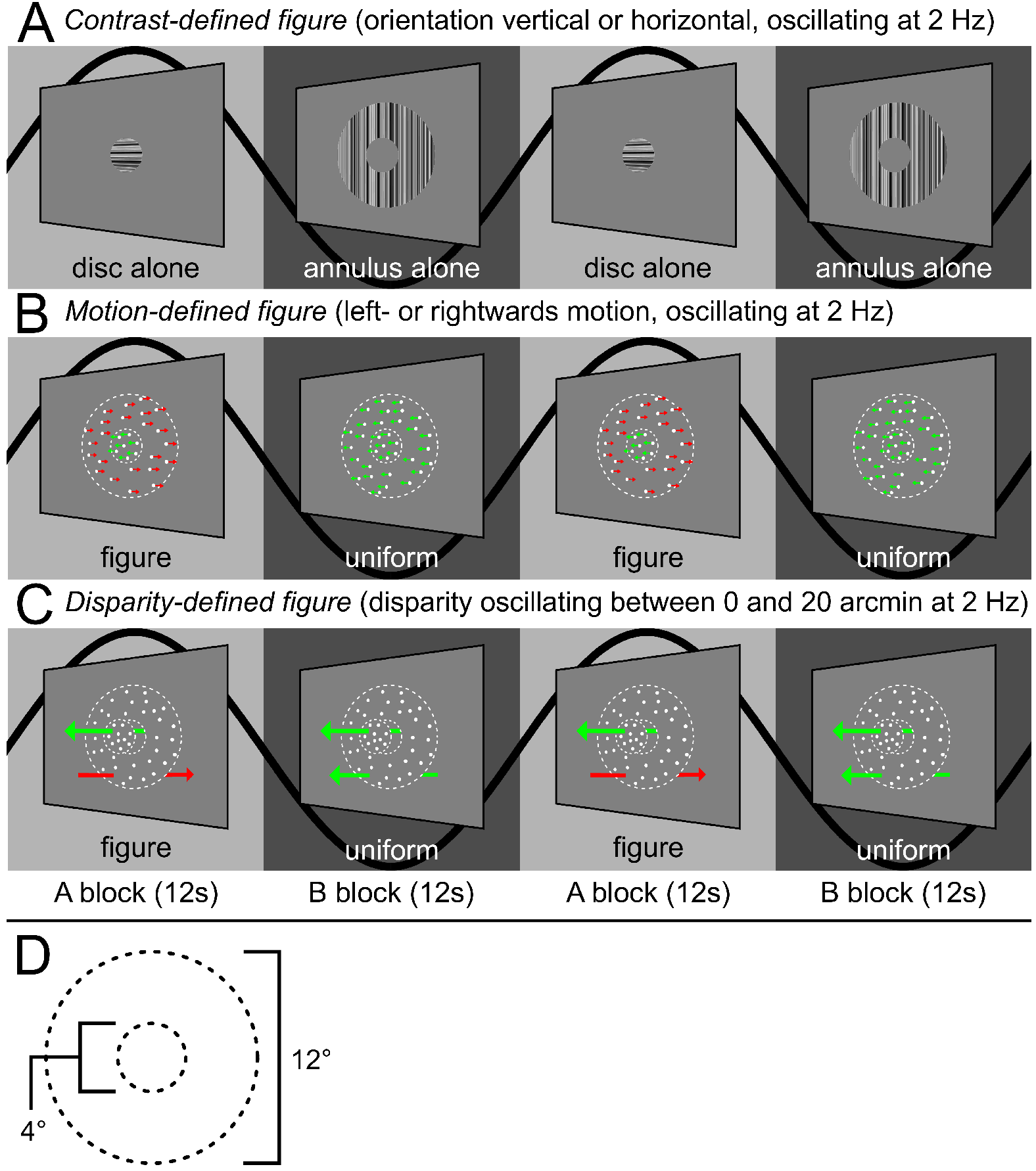
The experimental design used in Experiment 1. The on-off responses evoked by the conditions that are captured by our Fourier analysis, are illustrated with sine waves. (A) The contrast condition – the orientation of both the disc and annulus alternated between vertical and horizontal at 2 Hz. Disk and annulus were presented in temporal succession. (B) The motion condition – dots alternated between left- and right-wards motion at 2Hz. When the disc and annulus regions alternated in anti-phase, a figure percept was evoked, when alternations were in-phase, there was no such percept. (C) The disparity condition – the dots alternated between the fixation plane (0 arcmin) and a position behind fixation (uncrossed disparity, 20 arcmin) at 2 Hz. Anti-phase movement of the disc and annulus regions lead to a figure percept, while in-phase movement generated no such percept. The white dotted line in B and C indicates the extent of the disc and annulus used in A. Note that the relative disparity condition was identical to that used in Experiment 2, except for the introduction of an RSVP task at fixation, which directed attention away from the stimulus. (D) Schematic outlining the size of the disc and annulus region, respectively, which was shared across all conditions.

In areas of cortex defined by the template as responsive to eccentricities covered by the disc, we see a high amplitude disc responses. At eccentricities defined as near the disc boundary (2° radius), the phase sign reverses and we see annulus responses. There are significant responses at all eccentricities covered by the stimulus, except for the sign reversal region, where voxels responding to the disc and the annulus, 180° out of phase, are likely mixed together. The eccentricity of the sign reversal is slightly biased towards the periphery (about the size of the separation between sub-ROIs, 0.25°), but otherwise the pattern of results indicates that the template-based procedure allows us to accurately localize the eccentricity of the contrast-defined boundary in retinotopic cortex. The 180° phase reversal can also be directly observed in the whole-brain analysis of the contrast condition (Figure 6A), which plots the vector-averaged phase across all participants.

While the contrast condition alternated a disc and an annulus, the motion and disparity conditions alternated figure-ground and a uniform field configurations. The dots updated dynamically at 20 Hz throughout each block. In the disparity condition, they were temporally uncorrelated across updates, and thus did not generate monocular cues to form. In the motion case, the dots were temporally correlated with a long lifetime. The global structure of the displays (uniform vs segmented) updated at 2 Hz, alternating between left- and rightwards movement in the motion condition, and between the fixation plane (0 arcmin), and a position behind fixation (uncrossed disparity, 20 arcmin) in the disparity condition. The disc and annulus region alternated in anti-phase during the A block, and in phase during the B block. Importantly, absolute motion and absolute disparity were updating at 2 Hz during both A and B blocks, but only the A block generated relative motion and disparity cues that give rise to a figure boundary. The design and corresponding Fourier analysis yields the differential response and thus allows us to isolate responses driven by the presence of relative motion/disparity discontinuity, the segmented figure surface or some combination of the two from absolute motion/disparity responses.

The overall response amplitudes in the motion and disparity conditions were 5-10 times weaker than in the contrast condition, but there were nonetheless significant activations at multiple eccentricities. For both motion and disparity, positive amplitudes indicate stronger responses to the A block in which the figure and surround were
segmented (“figure responses”), while negative amplitudes indicate stronger responses to the B block in which a uniform field was perceived (“uniform responses”). For motion (see Figure 2B), we saw figure responses at the disc-annulus boundary, which persisted beyond the extent of the disc by 1° in V1 and V2, and even further in V3. There were also figure responses inside the disc region, which were most prominent in V2 and V3. This result indicates that we can detect responses to a figure boundary defined by relative motion in all three early visual areas. Surprisingly, we also saw uniform responses at eccentricities > 4.5°, which were significant in all visual areas, although less consistently in V1. This results indicates that when the boundary is present, responses in the region outside the boundary are weaker than when the boundary is not present. This is likely due to suppression of the surround when the boundary is present, rather than surround-exclusive enhancement of responses when the field is uniform. The lack of consistent suppression in V1 may be due to insufficient sensitivity of our method.

**Figure 2:**
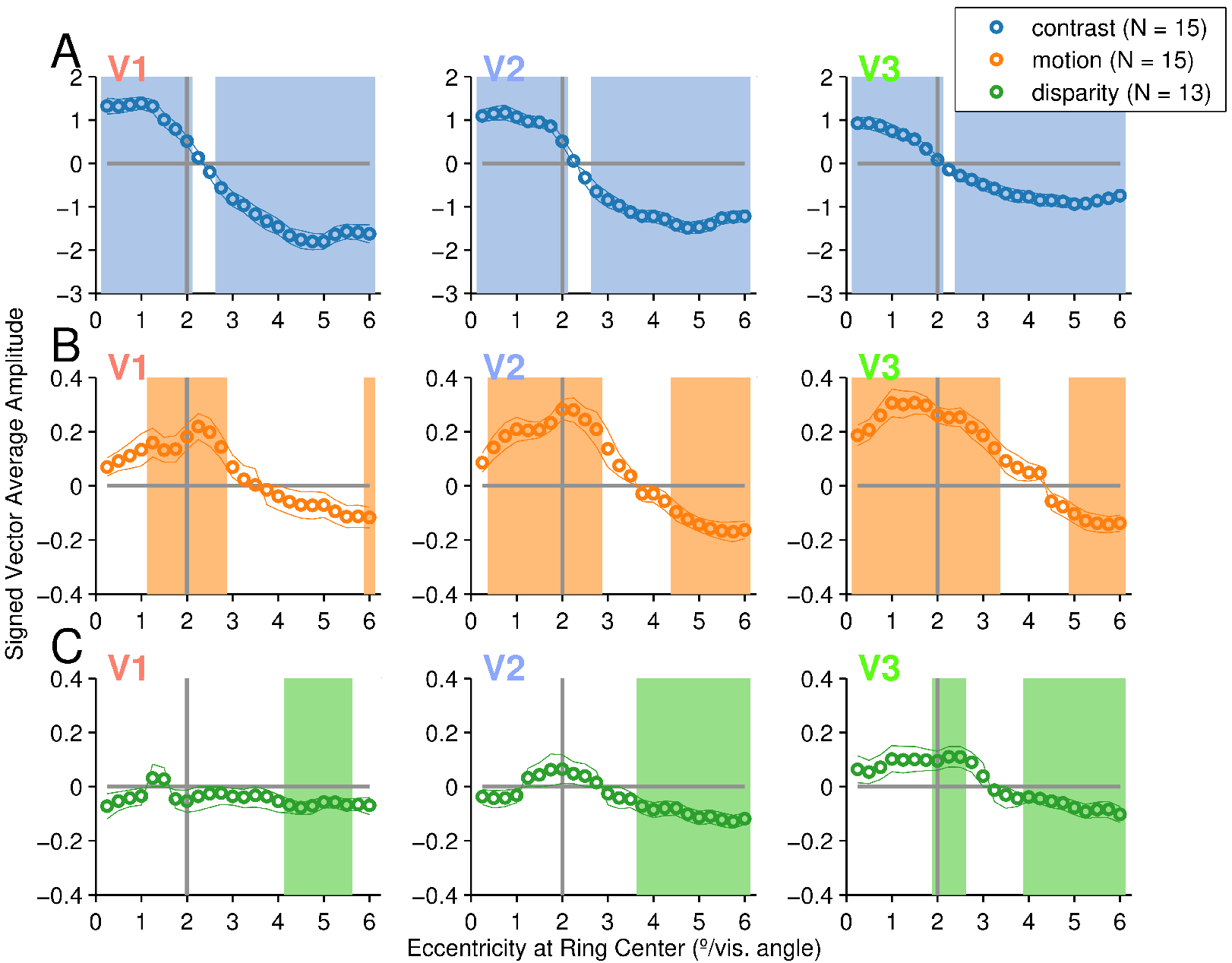
Eccentricity analysis of V1-V3 in Experiment 1. Signed vector mean amplitudes for the contrast (A), motion (B) and disparity (C) conditions within 24 eccentricity-defined sub-ROIs, centered on eccentricities spaced 0.25° apart, each having a width of 0.5°. The shaded areas on the plots indicate condition **×** sub-ROI combinations that were significant at α = 0.05. The color of the text for the ROI names matches the ROI colors on Figures 4 and 6.

In the disparity condition (see Figure 2C), we did not measure significant figure responses in V1 or V2 – although V2 responses at the boundary were positively signed, they were clearly not significant (lowest *p* = 0.445). In V3 by contrast, there were significant figure responses starting at the boundary and extending to ~0.5° outside the boundary. We also saw significant uniform responses in all early visual areas, which, as for the motion condition, are consistent with suppression that occurs during the A block when the boundary is present, but not in the B block when the field is uniform.

These results suggest an interesting dichotomy: Among the early visual areas, we only see evidence for responses to a figure boundary defined by relative disparity in V3, while all three early visual areas have evidence of suppression when the disparity-defined boundary is present. Relative suppression of surround-region responses may be due to feedback from higher cortical areas, driven by top-down attentional selection. Spatial attention can modulate BOLD responses in all early visual areas, including V1^38–40^. In Experiment 2, we directly tested the hypothesis that the suppression is due to attention-driven feedback.

Experiment 2 used the same parameters as the relative disparity condition in Experiment 1, except that attention was now directed away from the stimulus via an orthogonal task presented at fixation (see the ‘*Visual stimuli*’ section in the Methods). Under these conditions, we measured significant figure responses in V3, but not in V1 and V2, replicating results from Experiment 1 (see Figure 3). A single uniform response reached significance at the inner-most eccentricity in V2, but given the lack of spatial correspondence with the stimulus boundary and absence of any other trends in the V2 data, we consider this type 1 error. Importantly, we saw no evidence of a uniform response indicating suppression anywhere in any of the early visual cortex ROIs. This result support our hypothesis that the uniform response we saw in the surround region in Experiment 1 was due to top-down attentional suppression of the surround. Note that the figure-region response extended even further beyond the figure (~1.5°) than it did in Experiment 1 (~0.5°), perhaps due to a reduction of the negatively signed uniform responses in the surround. Our results also support the conclusion that the sensitivity of area V3 to a relative disparity-defined boundary that we measure as positively signed responses is not strongly dependent on attention.

**Figure 3:**
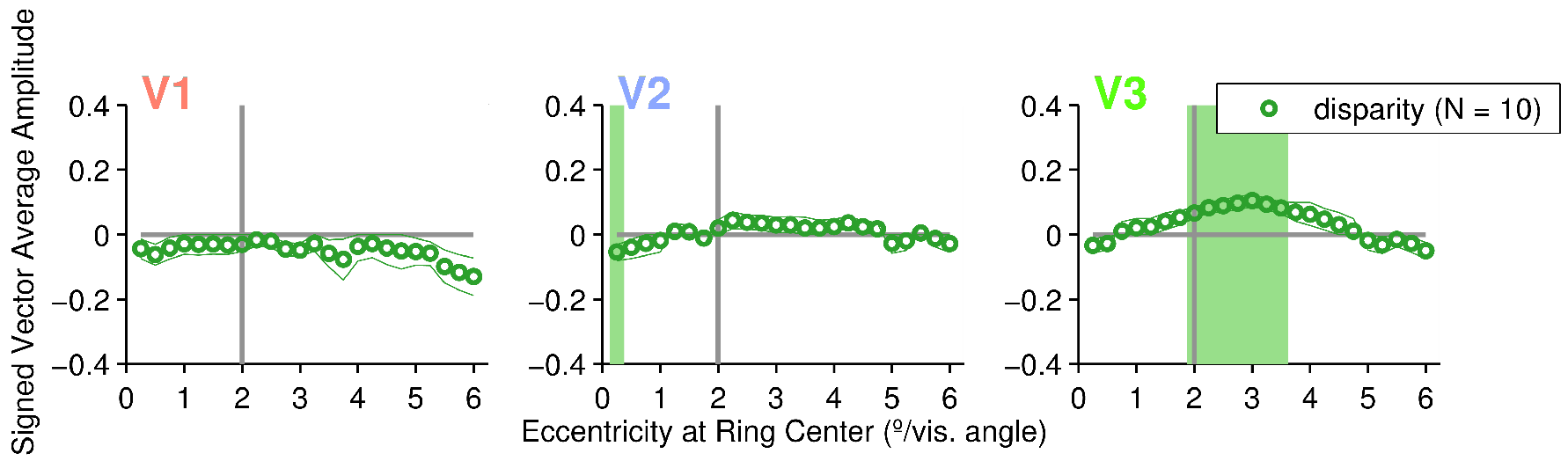
Eccentricity analysis of V1-V3 in Experiment 2. Signed vector mean amplitudes within 24 eccentricity-defined sub-ROIs, centered on eccentricities spaced 0.25° apart, each having a width of 0.5°. The shaded areas on the plots indicate condition **×** sub-ROI combinations that were significant at α = 0.05. The color of the text for the ROI names matches the ROI colors on Figures 4 and 6. The relative disparity condition shown here was identical to the one used in Experiment 1 (see Figure 2C), except attention was directed away from the stimulus.

### Extended ROI-based analysis of responsivity

The next set of analyses quantifies motion and disparity sensitivity in topographically organized visual areas beyond early visual cortex. Activation was defined as the signed vector mean amplitude of the average time course across all voxels within each ROI. The logic was the same as for the sub-ROI within early visual cortex: Positive amplitudes indicate responses to the figure (A block), while negative amplitudes indicate stronger responses to the uniform field (B block).

In both the motion- and disparity-defined form conditions, all significant activations were positively signed. This is consistent either with positive-sign activations generated by the figure overcoming any negative-sign activations that may have occurred in a subset of voxels within a given ROI, or a lack of negative-sign activation. We distinguished three response patterns: areas that only had significant responses (*p* < 0.05; indicated with shaded areas in Figure 4) to the motion-defined figure (hV4, TO1), areas that only had significant responses to the disparity-defined figure (PHC1, PHC2, IPS1, IPS2) and areas that had significant responses to both (V3B, LO1, LO2, IPS0, IPS3). Our results suggest a clear functional distinction between V3B and nearby area V3A, which has no significant responses. Motion generated stronger responses than disparity in V3 (see Figure 2), but this was less pronounced among the higher-level areas: Both V3B, IPS0 and IPS3 had nearly identical responses to the two conditions.

**Figure 4:**
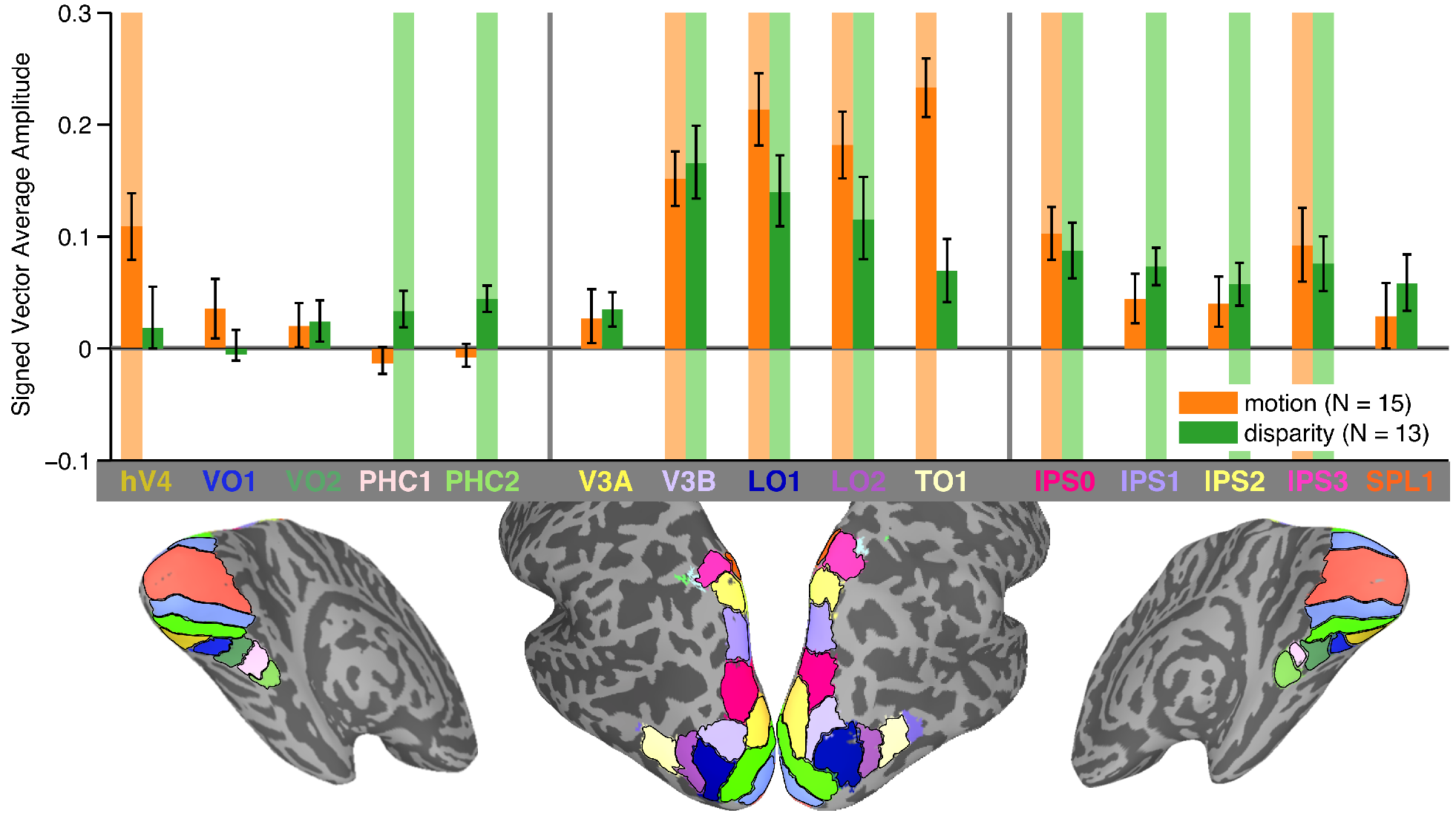
ROI results from Experiment 1. Signed vector mean amplitude within 15 topographically organized regions of interest, excluding early visual areas V1-V3, which are shown in Figure 2. The motion condition is shown in orange, and the disparity condition in green. Shaded areas behind the bars indicate condition × ROI combinations that were significant at α = 0.05. The ROIs are shown for the both hemispheres of an example participant’s inflated cortical surface reconstruction, below the graph.

### Effect of attention on disparity-defined figure activations outside of early visual cortex

We now ask if the areas that were sensitive to a disparity-defined figure boundary in Experiment 1, were also significant in Experiment 2, when attention was directed away from the stimulus via an orthogonal task at fixation. We found that responses persisted in areas V3B and LO1, but not in LO2 and IPS0-3 (see Figure 5), suggesting that activations in these latter areas depend on attention. Note that we were unable to probe PCH1 and 2 as these ROIs that were not covered by the fMRI acquisition protocol used in Experiment 2. hV4 also had a significant response (*p* = 0.008), which we did not see in Experiment 1 (*p* = 0.893). This difference in activation patterns could occur if hV4 has negative-sign, attention-dependent surround activations that cancel out the positive-sign figure activations. Our eccentricity analysis of early visual cortex demonstrated that these negatively-signed attention-driven effects are eliminated when top-down attention is controlled in Experiment 2. It is likely that the same thing is happening in hV4, eliminating the cancellation and revealing the positive-sign activations.

**Figure 5:**
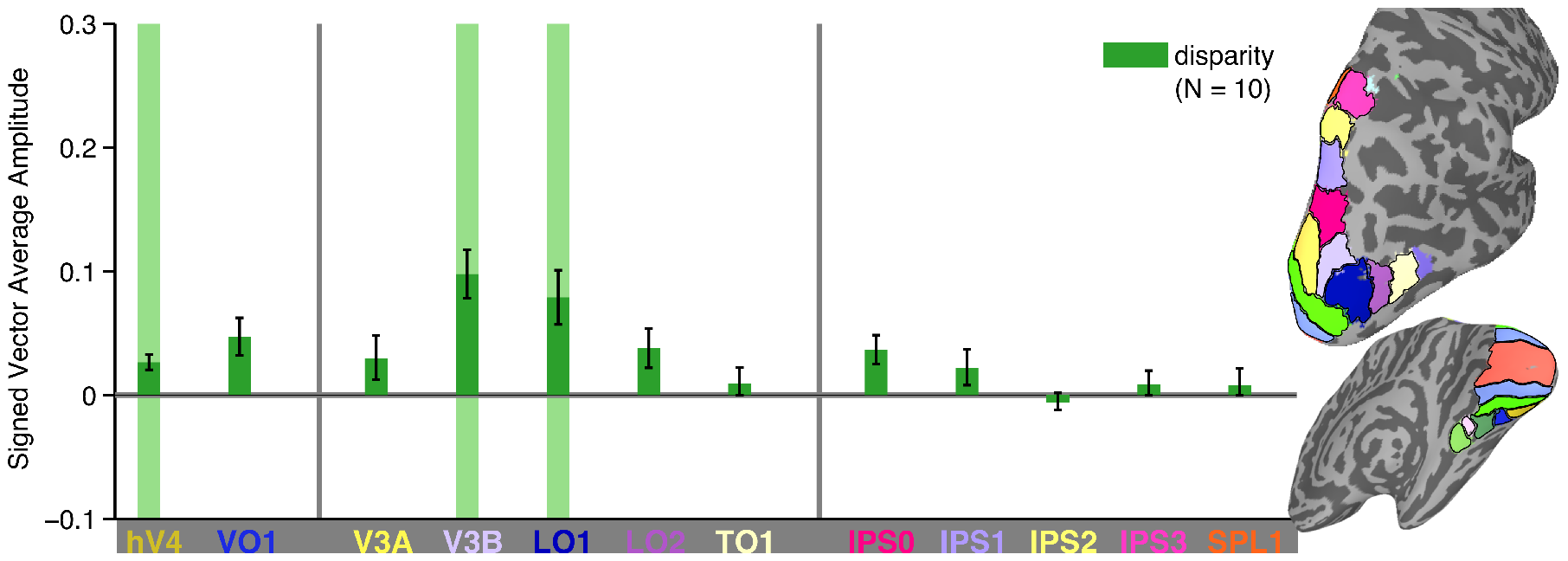
ROI results from Experiment 2. Signed vector mean amplitude within 12 topographically organized regions of interest, excluding early visual areas V1-V3, which are shown in Figure 3, and three ROIs that were not covered by the fMRI acquisition protocol used in Experiment 2 (VO2, PHC1 and PHC2). Shared areas behind the bars indicate condition × ROI combinations that were significant at α = 0.05. The ROIs are shown for the right hemisphere of an example participant’s inflated cortical surface reconstruction, on the right side of the graph.

### Whole brain analysis

A surface-based alignment approach was used to visualize vector-averaged responses to the conditions in Experiment 1, across all of cortex, including regions outside our set of ROIs. The results of these analyses were largely consistent with the sub-ROI and whole-ROI analyses, and we will only describe them briefly.

We plot the phase of the vector-averaged response, thresholded by significance. Blue colors indicate responses in phase with the A block, while orange colors indicate responses to the B block. For the contrast condition (see Figure 6A), the reversal of the phase sign from disc responses to annulus responses between low and high eccentricities, described in the sub-ROI analysis (see Figure 2A) can be clearly observed in early visual areas.

For motion, we see clear evidence in early visual cortex of both the figure region response at low eccentricities (Figure 6B, blue colors) and uniform responses consistent with the suppression of the surround (orange colors) that was described in the sub-ROI analysis. We also see figure responses that cover most dorsolateral ROIs and extend anteriorly and ventrally beyond the ROIs. In correspondence with the whole-ROI analysis (see Figure 4), we also see figure responses in IPS0, IPS3, and hV4, but not in V3A.

**Figure 6:**
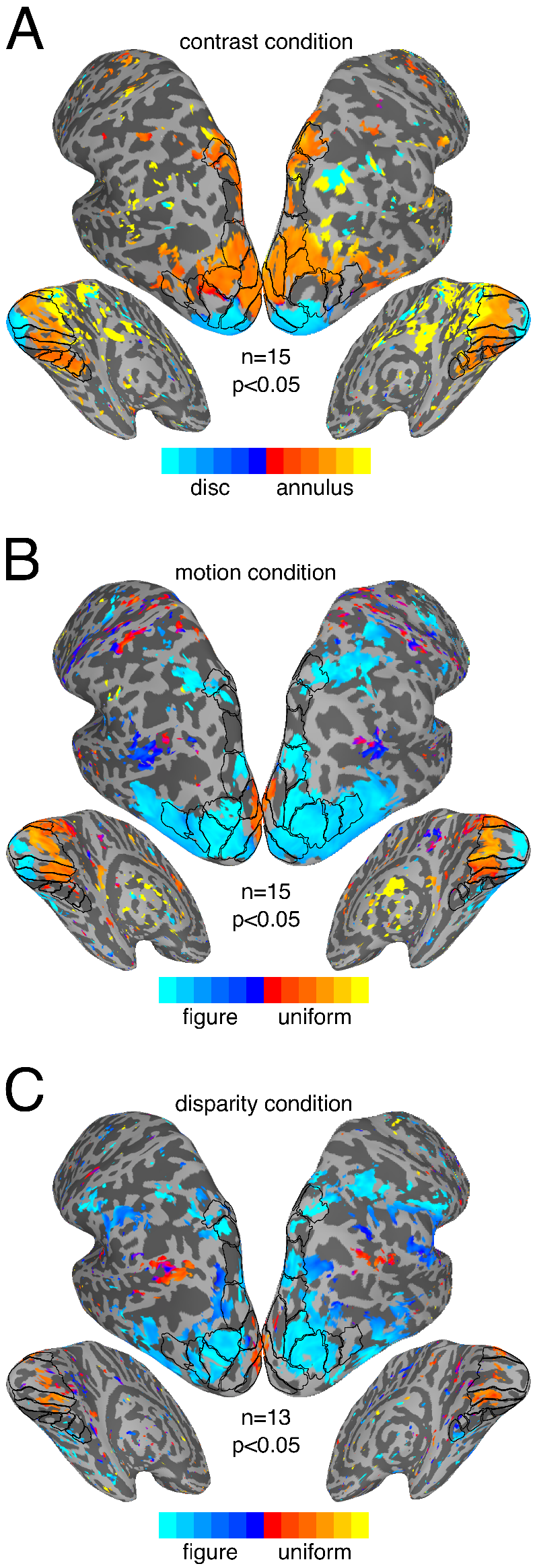
Whole-brain results from Experiment 1. Vector-averaged phase maps, thresholded at α = 0.05. Maps were produced through surface-based alignment procedure in which each subjects’ cortical mesh was converted to a standardized mesh, which allowed for cross-subject comparisons of values at each mesh node. The vector average phase across subjects, as well as a corresponding *p*-value based on both amplitude and phase, could then be computed for each mesh node. Blue shades indicate responses to the A block, while orange shades indicate responses to the B block. The ROIs are outlined on the surface, and labeled version can be inspected in Figure 4 and 5.

For disparity, we again see uniform responses consistent with suppression in early visual cortex (see Figure 6C, orange colors). It is worth noting that for both motion and disparity, there is little evidence of suppression outside of early visual cortex, at least within our ROIs. We do not see evidence of figure responses in V3, likely because the surface-based alignment approach is less sensitive than the sub-ROI analysis (see Figure 2). We do see figure responses in V3B, LO1, and LO2 and several IPS areas, but not in TO1 and V3A, in correspondence with the whole-ROI analysis (see Figure 4).

## 4. Discussion

We find evidence for representations of relative motion-defined figure boundaries in all early visual areas, and evidence for an analogous representation defined by relative disparity at least as early as V3, largely consistent with prior work in macaque^22,23^. In these areas, the activation patterns for both cues reflect the visual field topography of the stimulus, including a region of enhanced responses at or near the figure boundary, surrounded by suppressed responses in the ground region. We measure significant suppression associated with relative disparity in all early visual areas, but enhancement only reaches significance in V3, which may be due to lack of sensitivity of our method.

This pattern of results suggests that early visual areas go beyond simple edge-detection as reported previously in macaque V2^23^, and maintain representations related to the perceptual organization of the stimulus into figure and ground regions, as has been suggested for disparity on the basis of other single-unit recordings in macaque V2^24^.

Outside of early visual areas, we found areas that are sensitive to relative motion (TO1), relative disparity (PHC1, PHC2, IPS1, IPS2) or both (V3, hV4, LO1, LO2, V3B, IPS0, IPS3). Areas that are jointly selective contain candidate substrates for integrating these two cues. Ban and colleagues^1^ found evidence of integration of motion and disparity in V3B/KO, but not in any of the other five areas that we find to be sensitive to both cues. In Experiment 2, when attention was directed away from the stimulus using a letter task at fixation, disparity tuning was eliminated in a subset of the areas (LO2, IPS0-3), indicating that in those areas, sensitivity to disparity depends on attention. In V3, hV4, V3B and LO1, disparity tuning was independent of attention.

We observed stronger responses to relative motion compared to relative disparity in early visual areas (see Figure 2), but this difference was less pronounced in higher-level areas. This result is consistent with the ‘single-cue’ classification accuracies reported by Ban and colleagues, which are greater for motion than disparity in early visual areas, but more comparable in higher-level areas^1^. In some areas, their ‘single cue’ classification accuracies are above-chance for stimuli that do not produce significant responses in our data, most prominently in V1, V2 (disparity) and V3A (motion and disparity). It may be that in those cases, the classifier is picking up tuning for absolute motion and disparity^1^, which was controlled in our experiment design. Thus, the pattern of results in our current study and the study by Ban and colleagues^1^ suggest that V3A is sensitive to absolute, but not relative, motion and disparity, while neighboring V3B is sensitive to relative motion and disparity.

Relative disparity processing has been associated with the “canonical macaque ventral stream” leading from V4 to IT^41–43^. We find hV4 to be responsive to relative disparity as well as relative motion cues, which is consistent with reports that macaque V4 is sensitive to both relative disparity^26,27,44^, and relative motion^45^. In the macaque ventral stream, areas down-stream of V4 include sub-divisions of IT, TEO and TE, that are responsive to relative disparity^46,47^ and to relative motion^14,48^. In our data, areas immediately downstream of hV4, VO1 and VO2, are responsive to neither cue, while lateral surface areas LO1 and LO2 are responsive to both. Thus, for motion- and disparity-defined contours, the functional homology is poor between human ventral surface areas and macaque ventral areas, but better between human lateral surface areas and macaque ventral areas. Two recent proposals divide human visual cortex into three (dorsal, ventral and lateral) rather than the two, canonical dorsal and ventral streams of the macaque^49,50^. Our data suggests that functional homology exists between human lateral areas and the macaque ventral areas in IT cortex. Human ventral surface areas may either lack clear homologues in the macaque, or the homologous areas remain to be discovered.

Activations in V1, V2 and V3 for motion- and disparity-defined figures each consisted of a region of positive-sign activation surrounded by a negative-sign activation. The negative-sign activation in the surround could arise from suppression of responses during the A block or enhanced responses during the B block. There are, however, several reasons to suspect that suppression is the main driver of our negative-sign activation. First, the center and surround region are indistinguishable in the uniform stimuli used in the B blocks, and it is thus unlikely that there would be exclusive enhancement of the surround that would manifest as a negative-sign activation. Second, data from both macaque electrophysiology and human fMRI suggests that the relatively negative activations we see in the ground-region reflect suppression of neural responses. An opposing sign organization has been observed previously for “first-order” contrast-defined stimuli by contrasting blank screen baseline activation level to that within and adjacent to the retinotopic location of high contrast patterns^51–53^. Single unit and local field potential recordings in the retinotopic and adjacent representations of spatially localized, high contrast stimuli indicate suppression of baseline neural responses in the regions that had negatively signed BOLD^53^. Similar suppression may be occurring with our “second-order” contrast stimuli.

The center-surround configuration we used lends itself to the detection of alternate-sign activations because eccentricity is mapped systematically on the cortical surface in the foveal confluence-region of early visual cortex^54^. A suggestion of negative-sign or “out-of-phase” BOLD activation was present in the disparity data of Parker and Bridge^55^, but their use of rotating wedge-shaped stimuli complicated its visualization and measurement. Negative-sign activation has also been observed with a display in which a second-order figure region (a bar) was defined by temporal transients^56^. Negative-sign activation was found adjacent to the retinotopic locus of the bar, but that study did not find enhanced activation within the retinotopic representation of the figure as indicated by the positive-sign activations we observed^57^. Reppas and colleagues^18^ observed positive-sign activation at the border of a motion-defined form, but found neither negative-sign activation in the ground region nor positive-sign activation within the figure region, as we see in our data. Next, we will consider how suppression in the ground region could be driven by feedback from higher visual areas.

In Experiment 2, participants performed a demanding task at fixation that diverted attention away from the changing disparity stimuli. Under these conditions, the negative-sign activation in the surround was unmeasurable in all three visual areas, while the positive-sign activation at the figure region in V3 persisted. This suggests that the suppression was driven by feedback related to top-down attention, which decreased when participants no longer attended the figure. BOLD responses can be modulated by attention in a spatially specific fashion as early as V1^38–40^. If the surround suppression we observe is in fact due to attention-related feedback, we would expect to see it throughout early visual cortex, even in areas that are not sensitive to a boundary defined by relative disparity. The fact that negative-sign activation is observable for relative motion stimuli even at 6 degrees of retinal eccentricity in V1 is consistent with feedback from higher-level visual areas, as population receptive fields at this eccentricity, measured using fMRI, are on the order of 2 degrees or less^58,59^. Finally, we note that disparity tuning in IPS areas also disappeared when attention was captured at fixation, suggesting that these areas may be sources of feedback to early visual areas.

A surprising aspect of our data is the apparent expansion of the enhanced responses associated with the figure-region for disparity and motion-defined figures. Here we propose an explanation for these results, based on the way that fMRI voxels sample the visual field. Population receptive fields (pRFs) reflect a summary of the receptive fields of neurons sampled by each fMRI voxel,^60^ and are already quite large at 2° (>0.5° in V1, larger in V2 and V3^59^). This means that sub-ROIs both inside and outside the boundary will contain a mixture of voxels with pRFs that overlap with the boundary, and voxels with pRFs that do not. In the contrast condition, when the B block drives the surround as much as the A block drives the center, we would expect the sign reversal to occur at the boundary. It in fact occurs just outside (see Figure 2A), a bias that is likely due to the fact pRFs increase in size towards the periphery. This means that it is more likely that pRFs centered outside the disc will overlap with the disc than it is that pRFs centered inside the disc will overlap with the annulus, resulting in a bias in disc responses towards the periphery.

For the motion and disparity conditions in Experiment 1, the A block is driving the boundary while the surround is suppressed, leading to a phase sign reversal. If the relative suppression of the surround is weaker than the relative enhancement of the boundary, the enhancement might overpower the suppression, both in the summation of responses making up each pRF and in the averaging of pRFs within a sub-ROI. This would lead significant boundary enhancement to ‘survive’ outside the boundary; which is what we see in the motion and disparity conditions (see Figure 2B and C). If the B block response is further weakened, the reversal should move further towards the periphery, which is what we see in the disparity condition of Experiment 2 (see Figure 3). This simple model explains the boundary responses observed outside the figure and cannot be ruled out by our data. It is worth noting that the responses inside the figure, observed most prominently for motion in V2 and V3, properly reflect true enhancement of the figure, rather than pRFs sampling the boundary. pRFs are smaller towards the fovea and therefore less likely to overlap with the boundary.

Figure-ground segmentation could be supported by several mechanisms acting separately or in concert, including, as noted by Likova and Tyler^56^, retinotopic enhancement of the borders of the figure, retinotopic enhancement of the figure surface, retinotopic suppression of the ground region, and a combination of figure enhancement and ground suppression. There is both psychophysical^61^ and fMRI^57^ evidence for “competition-mediated ground suppression” in the absence of figure enhancement. A similar pattern of isolated suppression in the ground region was found with temporally defined forms^56^. Evidence for spatially antagonistic facilitative/suppressive interactions in figure-ground segmentation comes from an EEG source-imaging study^36^ in which relative disparity displays analogous to the one in the current study (segmented vs uniform) were contrasted with displays in which the center moved in depth within an uncorrelated surround and the complement (surround moving in depth, center uncorrelated). The response patterns could be modeled by a multiplicative interaction between center and surround in which the center response was enhanced and the surround response was suppressed in proportion to the magnitude of relative disparity. Our data from disparity- and motion-defined figures, combined with the single-unit electrophysiology for texture-defined form^62^ suggests that spatially antagonistic interactions may be a general computational strategy used across multiple stimulus domains. Our results are also consistent with the suggestion that boundary detection circuits in early visual cortex provide a structure for attentional selection^63^.

## Methods

### Participants

15 healthy adult participants (5 female; mean age = 30.6±13.5) participated in Experiment 1 and 10 participated in Experiment 2 (3 female; mean age = 37.4±15.3), with 1 participant taking part in both experiments. Each participant had visual acuity that was better than +0.1 LogMar (20/25) in each eye as measured on a Bailey-Lovie chart and stereo-acuity of 40 seconds of arc or better on the RandDot stereoacuity test. The experiment began after the procedures of the study had been explained and the participant had given written informed consent. Experiment protocol and consent forms were approved by the Stanford University Institutional Review Board, and all methods were performed in accordance with the relevant guidelines and regulations.

### Visual Stimuli

The stimuli for Experiment 1 were shown on a 47" Resonance Technology LCD display and viewed through a mirror at a distance of 277 cm. This resulted in a presentation area of 12.1 × 21.2 °/visual angle, of which our stimuli occupied 12 × 12°. The screen resolution was 1024 × 768 pixels, 8-bit color depth and a refresh rate of 60 Hz. In the relative disparity condition of Experiment 1, the mean luminance was 2.17 cd/m^2^ and contrast was 60%, and stereoscopic stimuli were displayed using red/blue anaglyph glasses, which were worn throughout the experiment. In the other two conditions of Experiment 1, the mean luminance was 34.49 cd/m^2^ and contrast was 90%. In Experiment 2, the stimulus was viewed through Resonance Technology LCD goggles, but the display parameters otherwise matched those used in the relative disparity condition of Experiment 1.

For each stimulus condition, the display comprised a central 2° radius disk region and an immediately adjacent 6° radius annulus. In the relative motion condition, the disk-boundary was defined using a random-dot kinematogram. In the relative disparity condition, the disk and the annulus were defined using dynamic random-dot stereograms with no monocular cues. For the relative disparity display, dot size was 5 minutes of arc (arcmin) and dot density was 36 per (°/visual angle)^2^, while in the relative motion display, dot size was 10.4 arcmin and dot density was 10 per (°/visual angle)^2^. We also ran a boundary localizer condition in which the disk-annulus boundary was defined by a contrast difference in texture patterns comprised of 1-dimensional noise which alternated between horizontal and vertical orientations at 3 Hz. This condition allowed us to compare the boundary activations found in the motion and disparity conditions to the activations generated by a contrast-defined boundary to which all visual areas should be highly sensitive. This localizer also served to verify the accuracy of the retinotopy template^37^ we used.

In the contrast condition, the central disk was presented in what we will refer to as the “A block” of the fMRI design and alternated with the adjacent annulus configuration, presented during the “B block” (see Figure 1A). In the relative motion condition, the horizontal positions of individual dots comprising a random dot pattern updated at 20 Hz. Dots were displaced by 10 arcmin per update (3.33 °/sec) with a dot life-time of 100 video frames. The dots moved leftwards or rightwards, changing direction at 1 Hz. The direction of motion inside and outside the central disk could either be in anti-phase, leading to a spatially segmented percept with a visible boundary between the disc and annulus regions defined by relative motion, or in phase, leading to a uniform motion percept with no boundary. In the A block of the fMRI design, the display alternated in a square-wave fashion between uniform motion and segmented configurations at 1 Hz. In the B block, only uniform motion was shown. Locally, each part of the display contained dots that alternated between leftward and rightward motion, only the relative direction of motion over the disk and annulus regions differed between A and B blocks.

In the relative disparity condition, the positions of individual dots updated at 20 Hz such that the dot fields were binocularly correlated but temporally uncorrelated (no monocular cues). The horizontal disparity of the central disk and the annulus alternated at 2 Hz between 0 disparity and 20 arcmin of uncrossed disparity. In the A block, the disc and annulus alternated in anti-phase, generating a spatially segmented percept with a visible boundary between the disc and annulus regions defined by relative disparity. In the B block, the disc and annulus alternated in phase, leading to a uniform motion percept with no border. Thus, disparity modulated between 0 and 20 arcmin at all locations in both A and B blocks, with only the relative disparity over the disk and annulus regions differing between A and B blocks. Participants wore anaglyph glasses throughout the experiment, but the contrast and motion condition were identical in both eyes and thus effectively shown at 0 disparity.

In Experiment 2 we replicated the relative disparity condition from Experiment 1, but introduced a rapid serial visual presentation (RSVP) task at fixation that served to direct attention away from the stimulus. Subjects attended to a letter F, randomly oriented and superimposed on the center of the display. At random times during a block the F briefly turned into either a target letter, either L or T, followed by a new F that served as a mask. On each change, subjects had to indicate with a button-press whether the target letter was an L or a T. The target letter duration was adapted online using a staircase procedure to stabilize performance at a constant level (~80% correct) during both A and B blocks.

### fMRI Experimental Procedure

We used a block design in which 12 s A blocks alternated periodically with 12 s B blocks, yielding a 24 s base period for the paradigm that was repeated 10 times in what
we refer to as a “scan”. The design is illustrated schematically in Figure 1. Ten stimulus cycles were shown per scan, with an additional half-cycle (one 12 s control block) being shown in the beginning of the scan to allow the brain and the scanner to settle. The data collected during this “dummy” period were removed from the fMRI time series data before the data analysis. The disparity condition was not run for 2 out of 15 participants in Experiment 1 because of technical issues. We acquired 4 scans per condition for each participant in Experiment 1, except 3/15 participants for whom we acquired only 3 scans for one or more of the conditions. In Experiment 2, we acquired 5 scans per participants, except 2/10 for whom we only acquired 4 scans.

### Structural and functional MRI acquisition

Functional and structural MRI data were collected on a General Electric Discovery 750 (General Electric Healthcare) equipped with a 32-channel head coil (Nova Medical) at the Center for Cognitive and Neurobiological Imaging at Stanford University. For each participant, we acquired two whole-brain T1-weighted structural datasets (1.0 × 1.0 × 1.0 mm resolution, TE=2.5 ms, TR=6.6 ms, flip angle=12, FOV=256 × 256) that were used for tissue segmentation and registration with atlas-based ROIS and retinotopy template (see below). For Experiment 1, a multiplexed EPI sequence ^64^ was used which allowed for the collection of 60 horizontal slices (2.0 × 2.0 × 2.0 mm resolution, TE = 30 ms, TR = 2000 ms, flip angle=77, FOV=220 × 220), resulting in whole-brain coverage. In Experiment 2, a non-multiplexed EPI sequence was used, which limited the coverage to 32 coronal slices, positioned at an oblique angle to maximize coverage of occipital, ventral and parietal cortices. The sequence used in Experiment 2 was otherwise identical to the one used in Experiment 1.

### fMRI analysis

After removing the dummy TRs, the fMRI data was preprocessed in AFNI ^65^, which included the following steps: slice-time correction, motion registration (the third TR of the first scan was always used as base), scaling (each voxel was scaled to a mean of 100, and values were clipped at 200), and de-trending (removing components corresponding to the six motion registration parameters, as well as Legendre polynomials of order 0 (constant signal), 1 (linear drift) and 2).

The remainder of the analysis was performed in MATLAB. The time-course data were first averaged across the three scans for each condition, and then across the voxels within each visual region-of-interest (ROI). We then applied a Fourier transform to the average time-course for each ROI, omitted DC, multiplied the spectrum by 2 to get the single sided spectrum, and scaled by dividing with the number of samples in the time-course. We selected the complex value at the stimulus frequency (10 cycles per scan) for each participant, within each ROI, and used it for statistical analysis. For the whole-brain analysis (see below), we performed the same Fourier analysis on a voxel-by-voxel basis, without averaging across ROIs, which gave us a complex value at the stimulus frequency, for every voxel in each participant.

### Visual regions-of-interest

Topographically organized visual ROIs were derived from a probabilistic atlas^66^. The atlas ROIs, defined by retinotopic mapping, included 25 ROIs covering 22 visual areas in ~50 individual participants. The atlas first converts the surface data from each individual to surface-based standardized space, and then converts the surface data from each individual to surface-based standardized space^67^, finally assessing the likelihood, across participants, of any particular vector on the standardized surface belonging to a particular ROI^66^. The atlas was defined using a maximum probability approach, which considers a given vector as part of the set of ROIs if it is more often found within the set, than outside the set, across participants. If this is the case, the vector is then assigned the value of the most likely ROI, and if not, it is considered to be outside the set of ROIs. The maximum probability approach captures much of the overall structure of ROIs defined for individual subjects and generalizes well to novel participants that did not contribute to the atlas generation^66^.

We downloaded the atlas from http://scholar.princeton.edu/sites/default/files/napl/files/probatlas_v4.zip and converted the ROIs from standardized surface space to native surface space for each of our participants, using nearest-neighbor interpolation. We removed vertices that were more than 1 edge away from the main cluster of each ROI, to ensure that all ROIs consisted exclusively of contiguous vertices. This step eliminated small speckles, while having minimal effect on the overall structure and extent of the ROIs. We then created a version of the structural data set associated with the surface meshes that was registered to the experiment data, and used that to convert the ROIs from surface space to volume space, registering them to the experimental data. When multiple surface nodes were mapped to a single voxel, the most common value across those nodes were assigned to the voxel. Finally, the ROIs were resampled to match the resolution and extent of the experiment data. We excluded four ROIs from our analysis, IPS4 and 5, TO2 and FEF, because of their small size in the probabilistic atlas, and merged the dorsal and ventral segments of V1, V2 and V3. This gave us a total of 18 bilateral ROIs to analyze.

To derive an independent estimate of the response in regions of early visual cortex responding to different eccentricities in the visual field, we used a template developed by Benson and colleagues^37^ that accurately predicts the location and retinotopic organization of early visual areas V1-V3, using only the cortical anatomy. After transforming the template data to match the specific cortical topology of each participant, we converted the eccentricity data in the template from surface space to volume space, and registered and resampled to match the experimental data, using the same approach as for the Wang ROIs. When multiple surface nodes were mapped to a single voxel, the average eccentricity value across those nodes were assigned to the voxel. We could now sub-divide the ROIs in early visual cortex, for each participant, by selecting voxels within V1, V2 and V3 that were responsive to a given range of eccentricities. We generated 24 sub-ROIs for each early visual area, centered on radii ranging from 0.25 to 6.0°, separated by 0.25°, and each spanning 0.5°/vis. angle.

### Vector-based statistics

We computed the average phase and amplitude at the stimulus frequency using a vector-based approach, in which the real and imaginary part of the complex value was averaged separately across participants, and then combined so that vector mean amplitude and phase could be computed. Error bars were computed using a geometrical approach, in which a two-dimensional error ellipse is computed, which describes the standard error of the mean response amplitude. The upper and lower error bounds were computed as the longest and shortest vectors from the origin to the error ellipse a detailed describtion of this approach can be found in ^68^. All statistical tests for significance were run as Hotelling’s *t*^2^ tests of the null hypothesis that the two-dimensional data set containing the real and imaginary parts of the complex value at the stimulus frequency was equal to [0,0]^69^. Note that this vector-based approach means that both amplitude and phase, and their consistency across participants, contributes to our reported estimates of mean amplitude, error and statistical significance.

We computed the sign of the responses by doing a linear fit with zero intercept of the real and imaginary values associated with the contrast condition, averaged across participants, within each of the eccentricity-based sub-divisions of V1. The amplitude of the response to the contrast condition was high at most retinotopic locations, but response phase varied with eccentricity. Values that were to the left and below a line orthogonal to the fit line were given negative phase signs (weaker responses to the A block than B block), while values above and to the right were given positive phase signs (stronger responses to the A block than B block). The contrast-based fit was used for all conditions. We multiplied mean amplitude with phase sign in the ROI plots, to illustrate the phase of the ROIs response.

### Whole-brain analysis

To provide an overview of the effect of our conditions across the whole brain, and account for any potential effects outside our set of visual ROIs, we mapped the complex values at each voxel onto a standardized cortical surface, for each participant. This surface-based alignment offers several advantages over volume-based approaches to group analysis^70^, most importantly by considering the structure of cortical sulci and gyri, as opposed to Talairach registration and other types cross-subject normalization in volume-space, which is likely to blur activations across neighboring banks of a sulcus^71^. After surface-based alignment, we used the real and imaginary parts of the complex value at the stimulus frequency, at each surface node across participants, to compute mean amplitude and phase using the same vector-based approach applied to the ROI data. We also performed the Hotelling’s *t*^2^ test for significance as described above, for each node on the surface, and used that for thresholding the surface data.

## Acknowledgements

This research was supported by NIH grant EY018775.

### Author Contributions

PJK, BRC and AMN designed the study. PJK and BRC collected the data. PJK analyzed the data. PJK, BRC and AMN wrote the paper.

### Additional Information

The authors declare no competing financial interests.

## References

1 Ban, H., Preston, T. J., Meeson, A. & Welchman, A. E. The integration of motion and disparity cues to depth in dorsal visual cortex. Nature neuroscience 15, 636–643, doi:http://www.nature.com/neuro/journal/v15/n4/abs/nn.3046.html-supplementary-information (2012).

2 Bradshaw, M. F. & Rogers, B. J. The interaction of binocular disparity and motion parallax in the computation of depth. Vision research 36, 3457–3468 (1996).

3 Nawrot, M. & Blake, R. Neural integration of information specifying structure from stereopsis and motion. Science 244, 716–718, doi:10.1126/science.2717948 (1989).

4 Lamme, V. A. F., van Dijk, B. W. & Spekreijse, H. Contour from motion processing occurs in primary visual cortex. Nature 363, 541–543 (1993).

5 Cao, A. N. & Schiller, P. H. Neural responses to relative speed in the primary visual cortex of rhesus monkey. Visual neuroscience 20, 77–84, doi:10.1017/S0952523803201085 (2003).

6 Lui, L. L., Bourne, J. A. & Rosa, M. G. P. Single-unit responses to kinetic stimuli in New World monkey area V2: Physiological characteristics of cue-invariant neurones. Experimental Brain Research 162, 100–108, doi:10.1007/s00221-004-2113-9 (2005).

7 Shen, Z.-M., Xu, W.-F. & Li, C.-Y. Cue-invariant detection of centre–surround discontinuity by V1 neurons in awake macaque monkey. The Journal of Physiology 583, 581–592, doi:10.1113/jphysiol.2007.130294 (2007).

8 Yin, J. et al. Breaking cover: neural responses to slow and fast camouflage-breaking motion. Proceedings of the Royal Society B: Biological Sciences 282, doi:10.1098/rspb.2015.1182 (2015).

9 Chen, M. et al. An Orientation Map for Motion Boundaries in Macaque V2. Cerebral cortex 26, 279–287, doi:10.1093/cercor/bhu235 (2016).

10 Allman, J., Miezin, F. & McGuinness, E. Direction- and Velocity-Specific Responses from beyond the Classical Receptive Field in the Middle Temporal Visual Area (MT). Perception 14, 105–126, doi:doi:10.1068/p140105 (1985).

11 Marcar, V. L., Xiao, D. K., Raiguel, S. E., Maes, H. & Orban, G. A. Processing of kinetically defined boundaries in the cortical motion area MT of the macaque monkey. J Neurophysiol 74, 1258–1270 (1995).

12 Raiguel, S., Van Hulle, M. M., Xiao, D. K., Marcar, V. L. & Orban, G. A. Shape and spatial distribution of receptive fields and antagonistic motion surrounds in the middle temporal area (V5) of the macaque. The European journal of neuroscience 7, 2064–2082 (1995).

13 Xiao, D. K., Raiguel, S., Marcar, V. & Orban, G. A. The spatial distribution of the antagonistic surround of MT/V5 neurons. Cereb Cortex 7, 662–677 (1997).

14 Sary, G., Vogels, R., Kovacs, G. & Orban, G. A. Responses of monkey inferior temporal neurons to luminance-, motion-, and texture-defined gratings. Journal of neurophysiology 73, 1341–1354 (1995).

15 Marcar, V. L., Raiguel, S. E., Xiao, D. & Orban, G. A. Processing of Kinetically Defined Boundaries in Areas V1 and V2 of the Macaque Monkey. Journal of neurophysiology 84, 2786–2798 (2000).

16 Dupont, P. et al. The kinetic occipital region in human visual cortex. Cereb Cortex 7, 283–292 (1997).

17 Van Oostende, S., Sunaert, S., Van Hecke, P., Marchal, G. & Orban, G. A. The kinetic occipital (KO) region in man: an fMRI study. Cerebral cortex 7, 690–701 (1997).

18 Reppas, J. B., Niyogi, S., Dale, A. M., Sereno, M. I. & Tootell, R. B. H. Representation of motion boundaries in retinotopic human visual cortical areas. Nature 388, 175–179 (1997).

19 Shulman, G. L., Schwarz, J., Miezin, F. M. & Petersen, S. E. Effect of Motion Contrast on Human Cortical Responses to Moving Stimuli. Journal of neurophysiology 79, 2794–2803 (1998).

20 Skiera, G., Petersen, D., Skalej, M. & Fahle, M. Correlates of figure-ground segregation in fMRI. Vision research 40, 2047–2056, doi:https://doi.org/10.1016/S0042-6989(00)00038-9 (2000).

21 Larsson, J., Heeger, D. J. & Landy, M. S. Orientation Selectivity of Motion-Boundary Responses in Human Visual Cortex. Journal of neurophysiology 104, 2940–2950, doi:10.1152/jn.00400.2010 (2010).

22 Thomas, O. M., Cumming, B. G. & Parker, A. J. A specialization for relative disparity in V2. Nature neuroscience 5, 472–478, doi:10.1038/nn837 (2002).

23 Bredfeldt, C. E. & Cumming, B. G. A simple account of cyclopean edge responses in macaque v2. J Neurosci 26, 7581–7596, doi:10.1523/JNEUROSCI.5308-05.2006 (2006).

24 Qiu, F. T. & von der Heydt, R. Figure and ground in the visual cortex: v2 combines stereoscopic cues with gestalt rules. Neuron 47, 155–166, doi:10.1016/j.neuron.2005.05.028 (2005).

25 Anzai, A., Chowdhury, S. A. & DeAngelis, G. C. Coding of Stereoscopic Depth Information in Visual Areas V3 and V3A. The Journal of Neuroscience 31, 10270–10282, doi:10.1523/jneurosci.5956-10.2011 (2011).

26 Umeda, K., Tanabe, S. & Fujita, I. Representation of Stereoscopic Depth Based on Relative Disparity in Macaque Area V4. Journal of neurophysiology 98, 241–252, doi:10.1152/jn.01336.2006 (2007).

27 Shiozaki, H. M., Tanabe, S., Doi, T. & Fujita, I. Neural Activity in Cortical Area V4 Underlies Fine Disparity Discrimination. The Journal of Neuroscience 32, 3830–3841, doi:10.1523/jneurosci.5083-11.2012 (2012).

28 Janssen, P., Vogels, R., Liu, Y. & Orban, G. A. Macaque Inferior Temporal Neurons Are Selective for Three-Dimensional Boundaries and Surfaces. The Journal of Neuroscience 21, 9419–9429 (2001).

29 Krug, K. & Parker, A. J. Neurons in Dorsal Visual Area V5/MT Signal Relative Disparity. The Journal of Neuroscience 31, 17892–17904, doi:10.1523/jneurosci.2658-11.2011 (2011).

30 Tsao, D. Y. et al. Stereopsis Activates V3A and Caudal Intraparietal Areas in Macaques and Humans. Neuron 39, 555–568, doi:https://doi.org/10.1016/S0896-6273(03)00459-8 (2003).

31 Backus, B. T., Fleet, D. J., Parker, A. J. & Heeger, D. J. Human Cortical Activity Correlates With Stereoscopic Depth Perception. Journal of neurophysiology 86, 2054–2068 (2001).

32 Mendola, J. D., Dale, A. M., Fischl, B., Liu, A. K. & Tootell, R. B. H. The Representation of Illusory and Real Contours in Human Cortical Visual Areas Revealed by Functional Magnetic Resonance Imaging. The Journal of Neuroscience 19, 8560–8572 (1999).

33 Neri, P., Bridge, H. & Heeger, D. J. Stereoscopic Processing of Absolute and Relative Disparity in Human Visual Cortex. Journal of neurophysiology 92, 1880–1891, doi:10.1152/jn.01042.2003 (2004).

34 Vinberg, J. & Grill-Spector, K. Representation of Shapes, Edges, and Surfaces Across Multiple Cues in the Human Visual Cortex. Journal of neurophysiology 99, 1380–1393, doi:10.1152/jn.01223.2007 (2008).

35 Cottereau, B. R., McKee, S. P., Ales, J. M. & Norcia, A. M. Disparity-Tuned Population Responses from Human Visual Cortex. The Journal of Neuroscience 31, 954–965, doi:10.1523/jneurosci.3795-10.2011 (2011).

36 Cottereau, B. R., McKee, S. P., Ales, J. M. & Norcia, A. M. Disparity-Specific Spatial Interactions: Evidence from EEG Source Imaging. The Journal of Neuroscience 32, 826–840, doi:10.1523/jneurosci.2709-11.2012 (2012).

37 Benson, N. C., Butt, O. H., Brainard, D. H. & Aguirre, G. K. Correction of Distortion in Flattened Representations of the Cortical Surface Allows Prediction of V1-V3 Functional Organization from Anatomy. PLoS computational biology 10, e1003538, doi:10.1371/journal.pcbi.1003538 (2014).

38 Gandhi, S. P., Heeger, D. J. & Boynton, G. M. Spatial attention affects brain activity in human primary visual cortex. Proceedings of the National Academy of Sciences 96, 3314–3319 (1999).

39 Martinez, A. et al. Involvement of striate and extrastriate visual cortical areas in spatial attention. Nature neuroscience 2, 364–369 (1999).

40 Somers, D. C., Dale, A. M., Seiffert, A. E. & Tootell, R. B. Functional MRI reveals spatially specific attentional modulation in human primary visual cortex. Proceedings of the National Academy of Sciences 96, 1663–1668 (1999).

41 Parker, A. J. Binocular depth perception and the cerebral cortex. Nature reviews 8, 379–391 (2007).

42 Neri, P. A stereoscopic look at visual cortex. J Neurophysiol 93, 1823–1826 (2005).

43 Verhoef, B. E., Vogels, R. & Janssen, P. Binocular depth processing in the ventral visual pathway. Philosophical transactions of the Royal Society of London 371, doi:10.1098/rstb.2015.0259 (2016).

44 Tanabe, S., Umeda, K. & Fujita, I. Rejection of false matches for binocular correspondence in macaque visual cortical area V4. J Neurosci 24, 8170–8180 (2004).

45 Mysore, S. G., Vogels, R., Raiguel, S. E. & Orban, G. A. Processing of Kinetic Boundaries in Macaque V4. Journal of neurophysiology 95, 1864–1880, doi:10.1152/jn.00627.2005 (2006).

46 Uka, T., Tanabe, S., Watanabe, M. & Fujita, I. Neural correlates of fine depth discrimination in monkey inferior temporal cortex. J Neurosci 25, 10796–10802 (2005).

47 Verhoef, B. E., Vogels, R. & Janssen, P. Inferotemporal cortex subserves three-dimensional structure categorization. Neuron 73, 171–182, doi:10.1016/j.neuron.2011.10.031 (2012).

48 Unno, S., Handa, T., Nagasaka, Y., Inoue, M. & Mikami, A. Modulation of neuronal activity with cue-invariant shape discrimination in the primate superior temporal sulcus. Neuroscience 268, 221–235, doi:10.1016/j.neuroscience.2014.03.024 (2014).

49 Haak, K. V. & Beckmann, C. F. Objective analysis of the topological organization of the human cortical visual connectome suggests three visual pathways. Cortex; a journal devoted to the study of the nervous system and behavior, doi:10.1016/j.cortex.2017.03.020 (2017).

50 Weiner, K. S. & Grill-Spector, K. Neural representations of faces and limbs neighbor in human high-level visual cortex: evidence for a new organization principle. Psychol Res 77, 74–97, doi:10.1007/s00426-011-0392-x (2013).

51 Saad, Z. S., DeYoe, E. A. & Ropella, K. M. Estimation of FMRI response delays. NeuroImage 18, 494–504 (2003).

52 Shmuel, A. et al. Sustained negative BOLD, blood flow and oxygen consumption response and its coupling to the positive response in the human brain. Neuron 36, 1195–1210 (2002).

53 Shmuel, A., Augath, M., Oeltermann, A. & Logothetis, N. K. Negative functional MRI response correlates with decreases in neuronal activity in monkey visual area V1. Nature neuroscience 9, 569–577, doi:10.1038/nn1675 (2006).

54 Dougherty, R. F. et al. Visual field representations and locations of visual areas V1/2/3 in human visual cortex. Journal of vision 3, 586–598, doi:10:1167/3.10.1 (2003).

55 Bridge, H. & Parker, A. J. Topographical representation of binocular depth in the human visual cortex using fMRI. Journal of vision 7, 15–15, doi:10.1167/7.14.15 (2007).

56 Likova, L. T. & Tyler, C. W. Occipital network for figure/ground organization. Experimental Brain Research 189, 257, doi:10.1007/s00221-008-1417-6 (2008).

57 Cacciamani, L., Scalf, P. E. & Peterson, M. A. Neural evidence for competition-mediated suppression in the perception of a single object. Cortex; a journal devoted to the study of the nervous system and behavior 72, 124–139, doi:10.1016/j.cortex.2015.05.018 (2015).

58 Amano, K., Wandell, B. A. & Dumoulin, S. O. Visual Field Maps, Population Receptive Field Sizes, and Visual Field Coverage in the Human MT+ Complex. Journal of neurophysiology 102, 2704–2718, doi:10.1152/jn.00102.2009 (2009).

59 Harvey, B. M. & Dumoulin, S. O. The relationship between cortical magnification factor and population receptive field size in human visual cortex: constancies in cortical architecture. The Journal of neuroscience: the official journal of the Society for Neuroscience 31, 13604–13612, doi:10.1523/JNEUROSCI.2572-11.2011 (2011).

60 Dumoulin, S. O. & Wandell, B. A. Population receptive field estimates in human visual cortex. NeuroImage 39, 647–660, doi:10.1016/j.neuroimage.2007.09.034 (2008).

61 Salvagio, E., Cacciamani, L. & Peterson, M. A. Competition-strength-dependent ground suppression in figure-ground perception. Attention, perception & psychophysics 74, 964–978, doi:10.3758/s13414-012-0280-5 (2012).

62 Poort, J., Self, M. W., van Vugt, B., Malkki, H. & Roelfsema, P. R. Texture Segregation Causes Early Figure Enhancement and Later Ground Suppression in Areas V1 and V4 of Visual Cortex. Cerebral cortex 26, 3964–3976, doi:10.1093/cercor/bhw235 (2016).

63 Qiu, F. T., Sugihara, T. & von der Heydt, R. Figure-ground mechanisms provide structure for selective attention. Nature neuroscience 10, 1492–1499 (2007).

64 Feinberg, D. A. et al. Multiplexed echo planar imaging for sub-second whole brain FMRI and fast diffusion imaging. PloS one 5, e15710, doi:10.1371/journal.pone.0015710 (2010).

65 Cox, R. W. AFNI: software for analysis and visualization of functional magnetic resonance neuroimages. Computers and Biomedical research 29, 162–173 (1996).

66 Wang, H. X., Merriam, E. P., Freeman, J. & Heeger, D. J. Motion Direction Biases and Decoding in Human Visual Cortex. The Journal of Neuroscience 34, 12601–12615, doi:10.1523/jneurosci.1034-14.2014 (2014).

67 Argall, B. D., Saad, Z. S. & Beauchamp, M. S. Simplified intersubject averaging on the cortical surface using SUMA. Human brain mapping 27, 14–27, doi:10.1002/hbm.20158 (2006).

68 Pei, F., Baldassi, S., Tsai, J. J., Gerhard, H. E. & Norcia, A. M. Development of contrast normalization mechanisms during childhood and adolescence. Vision research 133, 12–20, doi:https://doi.org/10.1016/j.visres.2016.03.010 (2017).

69 Anderson, T. W. An introduction to multivariate statistical analysis. (Wiley, 1984).

70 Hagler, D. J., Saygin, A. P. & Sereno, M. I. Smoothing and cluster thresholding for cortical surface-based group analysis of fMRI data. NeuroImage 33, 1093–1103, doi:https://doi.org/10.1016/j.neuroimage.2006.07.036 (2006).

71 Fischl, B., Sereno, M. I. & Dale, A. M. Cortical surface-based analysis: II: Inflation, flattening, and a surface-based coordinate system. NeuroImage 9, 195–207 (1999).

